# Age-related Inhibitory Decline: Examining Inhibition Sub-Components and their Impact on Sustained Attention in Healthy Ageing

**DOI:** 10.1101/2025.02.19.638971

**Authors:** Ciara Treacy, Sophie C. Andrews, Jacob M. Levenstein

**Author notes:** Correspondence concerning this article should be addressed to Ciara Treacy, Thompson Institute, University of the Sunshine Coast, 12 Innovation Pathway, 4575, Birtinya, Qld, Australia. Data analysis/experiment code and materials are openly available at the project’s Open Science Framework page (https://osf.io/xze4d/?view_only=bb6c327dd851463e9745cb74736b3993). The data appearing in this manuscript was presented as a poster at the Australasian Cognitive Neuroscience Society conference held in Newcastle, Australia (November 2024). We have no conflicts of interest to disclose. This work was completed with support from a Research Training Program Scholarship (University of the Sunshine Coast). We extend our sincere thanks to the participants who volunteered their time to contribute to this research.

## Abstract

**Objective:** The inhibition deficit hypothesis postulates that inhibitory functioning declines with age, which negatively impacts other cognitive abilities. Yet still, the impact of healthy ageing on inhibitory functioning remains unclear, with the multifaceted nature of inhibition often an overlooked factor. Moreover, no prior study has empirically tested whether inhibitory sub-components explain differential age-effects in sustained attention - an open question that this work aims to address.

**Method:** In this cross-sectional study, we investigated the inhibition deficit hypothesis in a sample of eighty healthy older adults (mean age = 67.78 years, 44f). We utilised the PsyToolkit platform to administer three inhibition tasks (i.e., flanker, Stroop, and go/no-go), each targeting a distinct sub-component process, along with the Sustained Attention to Response Task (SART).

**Results:** Semi-partial correlations (rho) of the three inhibition sub-component measures (correcting for gender and education) with age resulted in significant positive relationships with task performance on the Stroop and go/no-go, such that older individuals had more pronounced Stroop effects and worse go/no-go accuracy, respectively. Lastly, go/no-go performance completely mediated the relationship between ageing and sustained attention performance, whilst Stroop effects partially mediated this association.

**Conclusion:** Age-related declines were observed in specific inhibition tasks, which speaks against a *general* inhibition deficit in healthy ageing and cautions against conflating these sub-components into a generalised concept of inhibition. The mediation findings demonstrate that inhibitory sub-components account for age-related declines in sustained attention, over and beyond ageing itself via an indirect path, representing an important cognitive domain to maintain throughout ageing.

**Key points:** *Question:* Are different inhibitory sub-components associated with age, and do they mediate age-related changes in sustained attention?

*Findings:* Different inhibition tasks were uncorrelated, demonstrating specific relationships with age and providing evidence that select inhibition tasks can explain differential age-effects in sustained attention.

*Importance:* This study lends support for the inhibition deficit hypothesis and underlines the importance of considering inhibitions’ multifaceted nature.

*Next steps:* Future research should adopt longitudinal designs to examine causal relationships

---

Inhibition represents a core executive function that underlies the ability to voluntarily suppress impulses toward environmental interference and ignore competing distractions, building resistance against task-irrelevant information (Diamond, 2013.). Inhibition is crucial for achieving goal-oriented behaviour and managing the demands of everyday life. As a result, the study of inhibitory functioning has garnered much consideration, with evidence suggesting inhibition may also influence additional cognitive functions (Zacks et al., 2000; Zanto & Gazzaley, 2014). The inhibitory deficit hypothesis postulates that inhibitory functioning declines with age, and negatively influences broader cognitive domains (Hasher & Campbell, 2020; Hasher & Zacks, 1988). Behavioural evidence supporting an age-related inhibitory decline is found across a variety of commonly used experimental tasks, such as, the flanker (Colcombe et al., 2005; Zhu et al., 2010), Stroop (Andrés et al., 2008; Fong et al., 2021) and go/no-go (Nielson et al., 2002; Rey-Mermet & Gade, 2018) tasks. However, a consensus on the influence of age on inhibitory functioning has not been adopted fully.

Several studies have demonstrated no specific age-related deficits in inhibition (for flanker, see Hsieh and Fang (2012) & Salthouse (2010); for Stroop, see Borella et al. (2009); for go/no-go, see Grandjean and Collette (2011) & Vallesi (2011)), with two meta-analytical reviews exemplifying these contradictory findings. Verhaeghen (2011) analysed eight tasks across 119 studies and failed to identify any specific age-related deficits in inhibitory control, concluding that suggested declines in older adults have been largely overstated, and that inhibition may only have a minor role in the decline of cognition with age. More recently, Rey-Mermet and Gade (2018) examined eleven tasks across 196 studies and identified age-related impairments in only a select number of tasks (i.e., go/no-go and stop-signal task), calling the general inhibition deficit hypothesis into question. However, an often overlooked factor is that inhibition is not unitary, as evidenced by the numerous theoretical taxonomies that have been proposed (Friedman & Miyake, 2004; Nigg, 2000; Pettigrew & Martin, 2014; Stahl et al., 2014; Tiego et al., 2018). For example, Rey-Mermet and Gade (2018) propose three sub-components of inhibition: the ability to inhibit i) distracting information, ii) response interference, and iii) prepotent responding. In this study, we adopt their framework, which provides a recent classification of inhibition grounded in a diverse set of task-based measures. Neglecting the different classifications of inhibition, assuming generalisability beyond a particular sub-component process, likely exacerbates the apparent lack of consensus regarding age effects.

Further complicating behavioural assessment of inhibition is the understanding that individuals, particularly older adults (Starns & Ratcliff, 2010), tend to behave cautiously and prudently on response tasks, a strategy coined as the speed-accuracy trade-off (SAT). When a SAT is present, longer response times relate to preserved accuracy, depicted by a negative correlation between reaction times and error rates (Heitz, 2014). This concept is not a new phenomenon (Wickelgren, 1977), nor is it restricted to the study of inhibition (Kardos et al., 2020), for example, SATs have also been evidenced in sustained attention (Helton et al., 2009; Vallesi et al., 2021). Thus, not accounting for SATs also contributes to inconsistent findings and/or spurious effects.

In the context of healthy ageing, the inhibition deficit hypothesis has support (Fong et al., 2021; Persad et al., 2002), and opposition (Nicosia & Balota, 2020), however, deliberate attempts to determine whether inhibition can explain differential age-effects in other cognitive domains remain scarce. In sustained attention, the influence of ageing on performance is also a matter of ongoing debate (Lufi et al., 2015; Robison et al., 2022; Treacy et al., 2024). *Attention* requires top-down, goal-driven regulatory processes to resist distractions (Sarter et al., 2001), and therefore, inhibitory principles may form the foundation of sustaining attentional performance (Demeter et al., 2011). Thus, it is plausible that an age-related decline in inhibition may impact sustained attention abilities. To our knowledge, no study has empirically tested whether the inhibition deficit hypothesis extends to the domain of sustained attention, which may help explain the age-related disparities also prevailing in research on sustained attention and ageing.

Here, we acquired three distinct measures of inhibition, administering a different, commonly utilised task to index each theoretical sub-component. Specifically, the letter flanker task was utilised to assess the inhibition of distracting information, the Stroop colour and word test was utilised to evaluate the inhibition of response interference, and finally, a visuomotor go/no-go task was utilised to assess one’s capacity to inhibit prepotent motor responses. We compute standard performance outcome measures related to inhibition (i.e., reaction time and error rate congruence scores for flanker and Stroop, error rate for go/no-go) and, where appropriate, we compute SAT scores. As a preliminary step, we examined bivariate relationships among the three tasks probing different sub-components of inhibition, to understand how these tasks relate. We then investigated the hypothesis of an age-related inhibitory deficit, predicting that age would be associated with worse performance on all three tasks of inhibition. Next, we examined whether measures of inhibition explain age-related performance deficits in sustained attention. Consistent with the inhibition deficit hypothesis, it was hypothesised that inhibition would mediate the effects of age on sustained attention performance.

## Method

### Transparency and Openness

This study was approved by the [university blinded for review] Human Research Ethics Committee ([approval number blinded for review]), and all participants provided written, informed consent prior to study commencement. The study design, hypotheses, and analysis plan were not preregistered. All data pre-processing and analyses were performed using R Statistical Software v4.3.2 (R Core Team, 2023). The data analysis/behavioural experiment code and materials are available at https://osf.io/xze4d/?view_only=bb6c327dd851463e9745cb74736b3993. Data sharing has not yet been approved by the [university blinded for review] Human Research Ethics Committee but the data can be made available upon request by contacting the corresponding author. We report how we determined our sample size, any data inclusions/exclusions, measures and manipulations.

### Participants

Healthy older adults aged 50-85 years were recruited from the general community through a study specific webpage, media announcements and local community groups. Eligible participants were right hand dominant, English-speaking, generally healthy individuals without a diagnosis of mild cognitive impairment, psychiatric disorders (e.g., schizophrenia or bipolar), or major neurological conditions (e.g., dementia or Parkinson’s). Additionally, participants were absent of magnetic resonance imaging (MRI) contraindications (e.g., pacemaker, stents), colour-blindness, respiratory conditions (e.g., COPD), cardiovascular conditions (e.g., stroke, myocardial infarction), prior head injuries (loss of consciousness >60mins), poorly controlled diabetes, current substance abuse or misuse, and usage of medications known to impact the central nervous system (e.g., anti-depressant or anti-anxiety medication). Eligibility criteria were operationalized via online self-report questionnaires and telephone interviews. To determine the sample size required for the primary age-behaviour correlation analysis, we referred to prior research utilising similar tasks to investigate inhibition (Andrés et al., 2008; Fong et al., 2021; Li et al., 2018; Sebastian et al., 2013). An a priori power analysis was conducted using G*Power software.

For a two-tailed bivariate model with *r* = 0.37, power = 0.80, and α = 0.05, a sample size of n=55 participants was required. For the subsequent mediation analysis, G*Power software was deemed inappropriate for estimating sample size. Instead, we followed the sample size guidelines provided by Fritz and Mackinnon (2007), who reported that a minimum sample size of n=71 is required to achieve power = 0.80 under the α =0.39, β = 0.39 condition using a bias-corrected bootstrap approach. In total, eighty-one individuals satisfied eligibility screening and were enrolled in this cross-sectional study. One individual was excluded post-hoc due to cognitive difficulties, thus, the final behavioural cohort consisted of eighty community-dwelling healthy older adults (M_age_ = 67.78 years, 44f) which was adequate to test the hypotheses (see Table 1 for descriptive statistics). Included participants were born in the following countries: 78.75% Australia, 12.5% England, 3.75% New Zealand, 1.25% Germany, 1.25% China, 1.25% Papua New Guinea, 1.25% Scotland. Of the Australian participants, 1.59% were of Aboriginal origin. Participants attended a single, 4-hour session at the [location blinded for review] to complete the study, with all data collection occurring between 30^th^ January 2023 and 20^th^ September 2023.

**Table 1.** Participant Sample Characteristics.

| **Characteristic** | **n** | **Mean(±SD)** | **Median** | **Min - Max** |
| --- | --- | --- | --- | --- |
| Age (years) | 80 | 67.78(9.64) | 70.00 | 50.00 – 84.00 |
| Gender (f, m) | 44, 36 | - | - | - |
| Education (years) | 80 | 16.16(3.76) | 15.50 | 9.00 – 26.00 |
| Flanker Effect (error rate) | 80 | 0.58(2.97) | 1.13 | -7.67 – 8.00 |
| Flanker Effect (reaction time) | 80 | 23.68(31.01) | 25.50 | -73.00 – 109.00 |
| Stroop Effect (error rate) | 80 | 1.89(4.04) | 0.00 | -3.33 – 16.00 |
| Stroop Effect (reaction time) | 80 | 89.12(63.62) | 79.00 | -11.50 – 290.00 |
| Go/no-go (error rate) | 79 | 9.49(8.34) | 6.67 | 0.00 – 34.33 |
| Go/no-go (bis) | 79 | 0.02(1.11) | -0.06 | -3.53 – 2.15 |
| SART ($d'$) | 78 | 1.40(0.42) | 1.38 | 0.36 – 2.52 |
| SART (bis) | 78 | 0.09(1.15) | 0.23 | -2.37 – 2.85 |

n (sample size); female(f); male(m); balanced integrated score (bis); Sustained Attention to Response Task (SART); discrimination index ( $d'$ ).

### Behavioural tasks

Participants were seated in a quiet room at arm’s length from a DELL laptop screen (15.6” Precision 5530 model, intel UHD graphics 630), which was used to administer the behavioural experiments. All behavioural data was acquired via the PsyToolkit platform v3.4.2 (Stoet, 2010, 2017). For each task, participants were directed to maintain finger contact with the response key/s for the duration to ensure accurate record of reaction times and detection of slips of action. Further, participants were also instructed to give equal importance to both the speed and accuracy of their responses. Participants could take a short break (∼ 5 minutes) in between each task before the next set of instructions were given. Behavioural task diagrams are presented in Figure 1.

**Figure 1.**
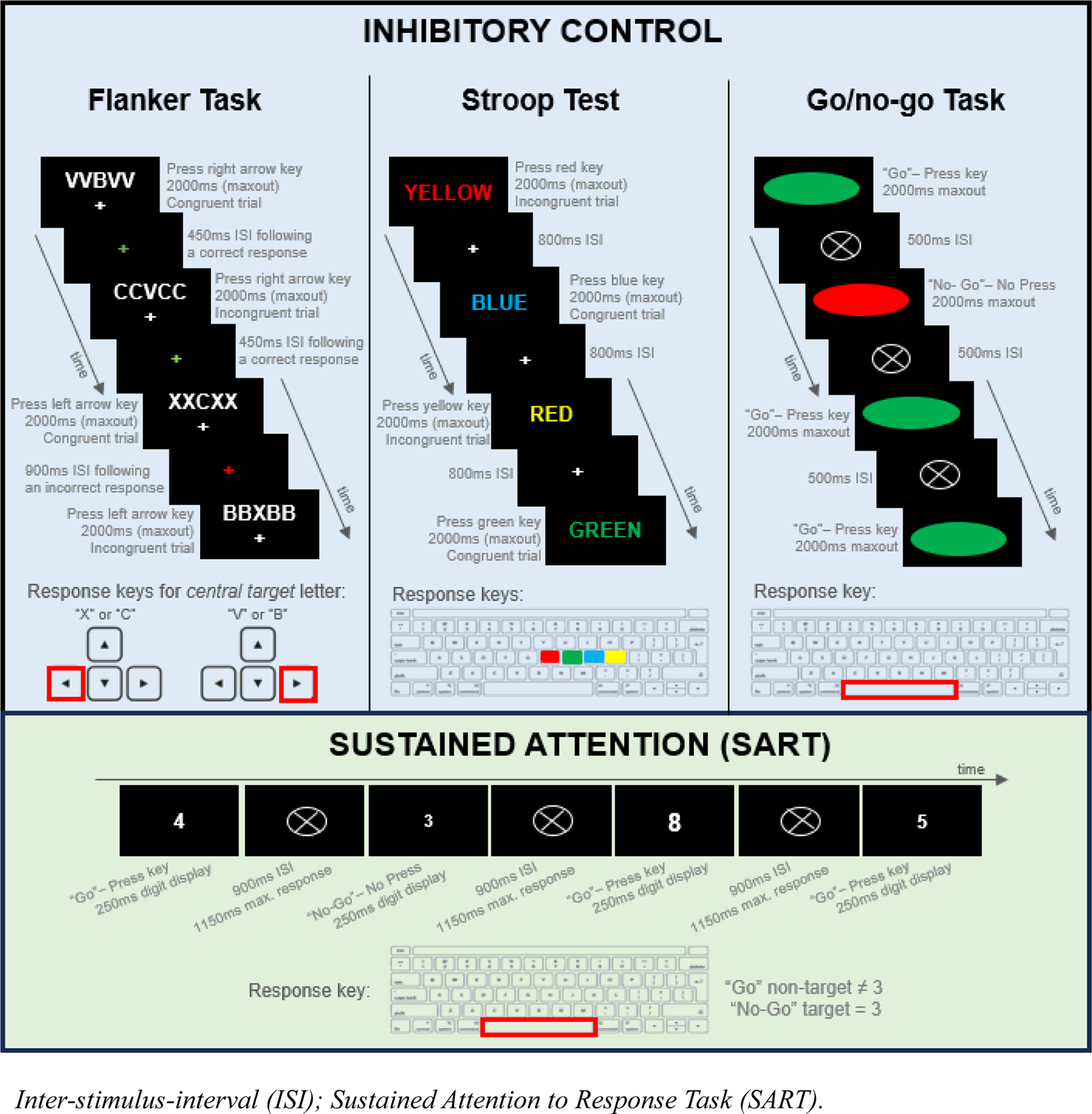
Behavioural Task Diagrams for Measures of Inhibition and Sustained Attention

### Flanker task

To assess distractor processing, participants completed a letter flanker paradigm (Eriksen & Eriksen, 1974), a non-search congruence task, involving 5 white coloured letters presented in a string above a white fixation cross in the centre of a black background (see Figure 1). Participants were instructed to register a response for the central letter only (i.e., the target), ignoring the two flanking distractor letters on either side (i.e., non-targets), by pressing the left arrow key if “X” or “C” were presented in the centre position, or, by pressing the right arrow key if “V” or “B” were presented in the centre position. Each 5-letter stimulus was displayed for 2000ms, which was also the maximum response window, followed by a fixation cross, incorporating performance feedback. More specifically, the inter-stimulus-interval (ISI) of the presented fixation cross varied depending on the prior response accuracy, such that correct responses presented a green fixation cross for 450ms, whereas incorrect responses were followed by a red fixation cross for 900ms. Participants completed a practice block (30 trials) before completing the 150 experimental trials. This task involved approximately equal proportions of congruent (47.3 – 52.7% of total) and incongruent (47.3 – 52.7% of total) trials, where congruent trials facilitate target responding, whilst response-incongruent trials incorporate visual conflict impacting target response execution. Flanker performance was measured by comparing congruent and incongruent trials in terms of median reaction times and error rates. Of note, we chose to include a congruence score for error rate to address a gap highlighted by Rey-Mermet and Gade’s (2018) meta-analysis, who found there was insufficient research to determine age-related deficits in flanker task accuracy. Flanker effect difference scores were calculated for each metric, such that flanker effect*_reaction time_* = incongruent_reaction time_ – congruent_reaction time_ and flanker effect_error rates_ = incongruent_error rates_ – congruent_error rates_. Flanker effects >0 reflect delayed and/or less accurate responding for incongruent trials, indicating that irrelevant flanking distractors (which are not compatible with the required response) influence task performance. Statistical summaries for the probability of reaction time and error rates for each trial type constituting the flanker effect scores are provided in the Supplemental Material (see Table S1).

### Stroop task

To assess inhibition of response interference, participants completed a Stroop paradigm (Golden, 1978), a colour-word interference task incorporating congruent and incongruent trials, involving one of four colour words (red, green, blue, yellow) being presented in the centre of a black background (see Figure 1). These colour words were presented in print colours which matched (i.e., response-congruent) or contradicted (i.e., response-incongruent) the meaning of the word itself. Participants were tasked to register a response relating only to the print colour (ignoring the colour word itself) by pressing the corresponding colour on the keyboard. Red, green, blue and yellow coloured stickers were placed over the h, j, k, and l keys, respectfully. Each colour-word stimulus was presented for 2000ms, which was also the maximum response window, followed by an 800ms white fixation cross (digit to digit onset was 2800ms maximum). Following a practice block of 16 trials, incorporating performance feedback, participants completed 120 experimental trials absent of performance feedback. This task involved equal proportions of congruent and incongruent trials, where congruent trials facilitated automatic responding, whilst incongruent trials introduced interference on automatic processing. In parallel with the flanker task, Stroop performance was also measured by comparing congruent and incongruent trials in terms of median reaction time and error rates, yielding Stroop effect difference scores for each metric. Likewise, we included a congruence score for errors in line with a recent meta-analysis that found more research was necessary to determine age-related deficits in Stroop task accuracy (Rey-Mermet & Gade, 2018). Stroop effects >0 reflect delayed and/or less accurate responding for incongruent trials, indicating that performing a less automated task, whilst inhibiting interference, influenced task performance. Statistical summaries for the probability of reaction time and error rates for each trial type constituting the Stroop effect scores are provided in the Supplemental Material (see Table S1).

### Go/no-go task

To assess the inhibition of prepotent responding, participants completed a go/no-go task, whereby a red or green coloured ellipse were presented one at a time in a pseudorandom order in the centre of a black background (see Figure 1). Each stimulus was presented for 2000ms, which was also the maximum response window, followed by a 500ms white fixation cross mask. Participants were instructed to register a response, by pressing the spacebar, for the green coloured ellipse (the “go” signal) and to withhold a response for the red coloured ellipse (the “no-go” signal). This task design involves frequent go events and rare no-go stimuli in a ratio of 4:1 (respectively), designed to build a prepotency to respond. Following a practice block of 25 trials, which incorporates performance feedback, participants completed 225 experimental trials (45 x no-go signals), absent of performance feedback. Trials with a response time greater than 700ms or less than 200ms were removed. Go/no-go performance was measured using *er*, determined as the proportion of commission errors (i.e., erroneous response to no-go stimuli), a metric frequently used as an index of response inhibition.

Additionally, a balanced integrated score (bis), which attenuates SAT’s, was calculated using the standardised mean difference between the proportion of correct responses and mean reaction times on correct trials, such that bis = z(x̅ percentage correct) - z(x̅ reaction time) (Liesefeld & Janczyk, 2019). The bis thereby represents a performance measure that controls for SAT’s and maintains “real” effects, with higher scores indicating better performance. Statistical summaries for reaction time and accuracy data constituting the bis density measure are also provided in the Supplemental Material (see Table S2).

### Sustained attention to response task (SART)

To assess sustained attention, participants completed the SART, which involves white coloured digits, ranging from 1 – 9, presented one at a time with equal probability in the centre of a black background in varying Arial font sizes (48, 72, 94, 100, 120), see Figure 1. Each digit was presented for 250ms followed by a 900ms mask (white fixation cross), with a maximum response window of 1150ms. Participants were asked to register a response, by pressing the spacebar, for all digits except 3, which represents the no-go target. Following a practice block of 18 trials, which incorporates performance feedback, participants completed 225 experimental trials (25 blocks of 9 trials), absent of feedback. For each block, a single digit was randomly chosen without repeating, therefore, over the course of the experimental phase no-go targets represent 11.1% of total trials. SART performance was measured using the discrimination index (*d’*), calculated using the standardised difference between the means of omission error (hit rate (h)) and commission error (false alarms (f)) distributions, such that *d’* = Z(h) – Z(f). The *d’* represents the ability to detect targets from non-targets, with higher scores indicating better signal detection/sustained attention performance. Log-linear correction was applied to the full dataset to address the issue in signal detection theory regarding participants with extreme values (i.e., h=1; f=0). Additionally, a bis was also calculated as previously described to provide a comparative measure which controls for SAT’s. Statistical summaries for the probability of omission and commission errors as well as reaction time data constituting the density performance measures (i.e., *d’* and bis) are provided in the Supplemental Material (see Table S3).

### Statistical Analyses

Outlier adjustment was performed as required using the z-score standard deviation transformation method, such that z-scores >3.29 or <3.29 were considered extreme cases and subsequently adjusted to one unit above or below the nearest value existing within acceptable ranges (Tabachnick et al., 2013). Three univariate outliers were adjusted in the Stroop data, and one univariate outlier was adjusted in the flanker and go/no-go data. Univariate analyses were performed to generate descriptive summaries. To provide an estimate of internal consistency of the behavioural measures, we used the splithalf package in R (version 0.8.2), which implements a permutation-based splithalf approach with 5000 random splits (Parsons, 2021; Parsons et al., 2019). Initially, bivariate correlations were performed across the three inhibition measures using Spearman’s rho (2-tailed). Then, semi-partial correlations between age and inhibition measures were performed, correcting behavioural performance metrics for gender and education to isolate age-effects. To account for multiple comparisons, we applied a Bonferroni correction resulting in a more conservative alpha for behaviour-behaviour correlations (8 tests) of p≤0.006 and age-behavioural correlations (6-tests) of p≤0.008. We also used the BayesFactor package in R (version 0.9.12-4.6) to calculate Bayes Factors (BF_10_) with a wide Cauchy prior distribution (Morey & Rouder, 2023) to assess the strength of evidence for the alternate hypothesis (H_1_) relative to the null hypothesis (H_0_). The BF in favour of H_0_ was also reported (i.e., BF_01_). Following Jeffrey’s (1961) guidelines, a BF between 1 and 3 is considered anecdotal evidence, 3 < BF <10 is considered moderate evidence, 10 < BF < 30 is considered strong evidence, 30 < BF < 100 is considered very strong evidence, and BF >100 is considered extreme evidence. Lastly, mediation analyses were performed using the PROCESS macro for R (version 4.3.1; mediation model 4 with a bootstrapping 5000; Preacher and Hayes (2008)). For each mediation model, gender and years of education were entered as covariates, age was entered as the independent variable, one inhibition measure per analysis was entered as the mediator, with a measure of sustained attention (SART *d’* or SART bis) representing the dependent variable in each instance.

Mediation effects were only tested for significant age-inhibition relationships resulting from the semi-partial correlations. Mediation was determined if the indirect effects were significant (i.e., *t-*values > 1.96 and the confidence intervals do not cross zero).

## Results

All participants completed the flanker and Stroop tasks, one individual did not complete the go/no-go task whilst two individuals did not complete the SART (Table 1). Thus, the following analyses were run for the complete datasets available, applying pairwise deletion. The Spearman-Brown corrected reliability estimate for Stroop effects (error rates) was r_SB_ = 0.80, Stroop effects (reaction time) was r_SB_ = 0.78, flanker effects (error rates) was r_SB_ = 0.30, flanker effect (reaction time) was r_SB_ = −0.14, go/no-go (error rates) was r_SB_ = 0.76, go/no-go (reaction time) was r_SB_ = 0.63, SART (error rates) was r_SB_ = 0.82, and SART (reaction time) was r_SB_ = 0.96 (see Supplemental Material, Table S4).

### Bivariate Relationships Among Inhibition Tasks

After controlling for multiple comparisons (adjusted p≤0.006), no significant correlations were identified between the flanker, Stroop or go/no-go measures (Table 2). Bayesian analysis of the correlation between flanker effects (reaction time) and go/no-go performance indicated moderate evidence in favour of the null hypothesis, both in terms of go/no-go errors (BF_01_ = 3.50) and bis scores (BF_01_ = 3.47, see Table 2). In contrast, correlations between Stroop effects (reaction time) and go/no-go performance were supported by moderate to strong Bayesian evidence for the alternate hypothesis, with BF_10_ = 6.50 for go/no-go errors and BF_10_ = 27.78 for bis scores (Table 3). Furthermore, for the correlation between Stroop effect (error rate) and go/no-go (bis), Bayes factors provided very strong evidence for the alternate hypothesis, BF_10_ = 61.57 (Table 2). For all remaining inter-task correlations, Bayes factors provided only anecdotal evidence (BF < 1, see Table 2).

**Table 2.** Bivariate Correlations (Spearman’s rho and Bayes factor) of Inhibition Measures.

|  | Flanker Effect<br>(error rate) | Flanker Effect<br>(reaction time) | Stroop Effect<br>(error rate) | Stroop Effect<br>(reaction time) | Go/no-go<br>(error rate) | Go/no-go<br>(bis) |
| --- | --- | --- | --- | --- | --- | --- |
| Flanker Effect<br>(error rate) | - | - | $\rho = -0.042$<br>$p_{raw} = 0.710$<br>$p_{cor} = 5.683$<br>$BF_{01} = 0.744$<br>$BF_{10} = 1.344$ | $\rho = -0.020$<br>$p_{raw} = 0.857$<br>$p_{cor} = 6.860$<br>$BF_{01} = 2.795$<br>$BF_{10} = 0.358$ | $\rho = -0.057$<br>$p_{raw} = 0.617$<br>$p_{cor} = 4.934$<br>$BF_{01} = 1.657$<br>$BF_{10} = 0.604$ | $\rho = 0.177$<br>$p_{raw} = 0.118$<br>$p_{cor} = 0.946$<br>$BF_{01} = 0.607$<br>$BF_{10} = 1.647$ |
| Flanker Effect<br>(reaction time) | - | - | $\rho = 0.160$<br>$p_{raw} = 0.158$<br>$p_{cor} = 1.264$<br>$BF_{01} = 1.624$<br>$BF_{10} = 0.616$ | $\rho = 0.005$<br>$p_{raw} = 0.966$<br>$p_{cor} = 7.731$<br>$BF_{01} = 2.422$<br>$BF_{10} = 0.413$ | $\rho = -0.102$<br>$p_{raw} = 0.370$<br>$p_{cor} = 5.116$<br>$BF_{01} = 3.498$<br>$BF_{10} = 0.286$ | $\rho = 0.051$<br>$p_{raw} = 0.654$<br>$p_{cor} = 5.233$<br>$BF_{01} = 3.470$<br>$BF_{10} = 0.288$ |
| Stroop Effect<br>(error rate) | $\rho = -0.042$<br>$p_{raw} = 0.710$<br>$p_{cor} = 5.683$<br>$BF_{01} = 0.744$<br>$BF_{10} = 1.344$ | $\rho = 0.160$<br>$p_{raw} = 0.158$<br>$p_{cor} = 1.264$<br>$BF_{01} = 1.624$<br>$BF_{10} = 0.616$ | - | - | $\rho = 0.042$<br>$p_{raw} = 0.714$<br>$p_{cor} = 5.710$<br>$BF_{01} = 0.649$<br>$BF_{10} = 1.541$ | $\rho = -0.282$<br>$p_{raw} = 0.012$<br>$p_{cor} = 0.095$<br>$BF_{01} = 0.016$<br>$BF_{10} = 61.565$ |
| Stroop Effect<br>(reaction time) | $\rho = -0.020$<br>$p_{raw} = 0.857$<br>$p_{cor} = 6.860$<br>$BF_{01} = 2.795$<br>$BF_{10} = 0.358$ | $\rho = 0.005$<br>$p_{raw} = 0.966$<br>$p_{cor} = 7.731$<br>$BF_{01} = 2.422$<br>$BF_{10} = 0.413$ | - | - | $\rho = 0.187$<br>$p_{raw} = 0.099$<br>$p_{cor} = 0.790$<br>$BF_{01} = 0.154$<br>$BF_{10} = 6.493$ | $\rho = 0.230$<br>$p_{raw} = 0.041$<br>$p_{cor} = 0.329$<br>$BF_{01} = 0.036$<br>$BF_{10} = 27.778$ |
| Go/no-go<br>(error rate) | $\rho = -0.057$<br>$p_{raw} = 0.617$<br>$p_{cor} = 4.934$<br>$BF_{01} = 1.657$<br>$BF_{10} = 0.604$ | $\rho = -0.102$<br>$p_{raw} = 0.370$<br>$p_{cor} = 5.116$<br>$BF_{01} = 3.498$<br>$BF_{10} = 0.286$ | $\rho = 0.042$<br>$p_{raw} = 0.714$<br>$p_{cor} = 5.710$<br>$BF_{01} = 0.649$<br>$BF_{10} = 1.541$ | $\rho = 0.187$<br>$p_{raw} = 0.099$<br>$p_{cor} = 0.790$<br>$BF_{01} = 0.154$<br>$BF_{10} = 6.493$ | - | - |
| Go/no-go<br>(bis) | $\rho = 0.177$<br>$p_{raw} = 0.118$<br>$p_{cor} = 0.946$<br>$BF_{01} = 0.607$<br>$BF_{10} = 1.647$ | $\rho = 0.051$<br>$p_{raw} = 0.654$<br>$p_{cor} = 5.233$<br>$BF_{01} = 3.470$<br>$BF_{10} = 0.288$ | $\rho = -0.282$<br>$p_{raw} = 0.012$<br>$p_{cor} = 0.095$<br>$BF_{01} = 0.016$<br>$BF_{10} = 61.565$ | $\rho = 0.230$<br>$p_{raw} = 0.041$<br>$p_{cor} = 0.329$<br>$BF_{01} = 0.036$<br>$BF_{10} = 27.778$ | - | - |

Balanced integrated score (*bis*). Bayes Factor in favour of $H_0$ ( $BF_{01}$ ) and $H_1$ ( $BF_{10}$ ). $p_{raw}$ = uncorrected $p$ value, $p_{cor}$ = corrected $p$ value.

**Table 3.** Semi-Partial Correlations (Spearman’s rho) between Age and Inhibition Measures.

|  | **Age (years)** |  |  |  |  |
| --- | --- | --- | --- | --- | --- |
|  | *Coefficient*<br>*(rho)* | *P-value*<br>*(raw)* | *P-Value*<br>*(corrected)* | *Bayes Factor* |  |
|  |  |  |  | *BF<sub>10</sub>* | *BF<sub>01</sub>* |
| Flanker Effect (error rate) | -0.030 | 0.792 | 1.000 | 0.329 | 3.044 |
| Flanker Effect (reaction time) | -0.051 | 0.654 | 1.000 | 0.326 | 3.072 |
| Stroop Effect (error rate) | 0.337 | **0.002** | **0.014** | 83.750 | 0.012 |
| Stroop Effect (reaction time) | 0.313 | **0.005** | **0.028** | 21.444 | 0.047 |
| Go/no-go (error rate) | 0.045 | 0.696 | 1.000 | 0.261 | 3.826 |
| Go/no-go (bis) | -0.471 | **<0.001** | **<0.001** | 391.630 | 0.003 |

Performance measures were residual corrected for gender and education; balanced integrated score (*bis*). Bayes Factor in favour of H<sub>1</sub> (BF<sub>10</sub>) and H<sub>0</sub> (BF<sub>01</sub>). Significant correlations (rho) are bolded.

### Age Relationships with Behavioural Inhibition

After correcting behavioural inhibition measures for gender and education, age was negatively correlated with go/no-go bis, such that higher age was associated with poorer overall go/no-go performance (see Table 3, Figure 2A). Bayesian analysis provided extreme evidence in support of the alternate hypothesis for this correlation, BF_10_ = 391.63 (Table 3). Additionally, age was positively correlated with Stroop effects in terms of reaction time (see Table 3, Figure 2B) and errors (see Table 3, Figure 2C), such that higher age was associated with slower and less accurate performance on the Stroop (more pronounced Stroop effect). These correlations were supported by strong to very strong Bayesian evidence for the alternate hypothesis, with BF_10_ = 83.7 for error rates and BF_10_ = 21.44 for reaction times (Table 3). No significant correlations were observed between age and flanker effects, or a traditional measure of go/no-go performance (i.e., commission errors), with Bayes Factors providing moderate evidence in favour of the null hypothesis for these comparisons (3<BF_01_<10, see Table 3).

**Figure 2.**
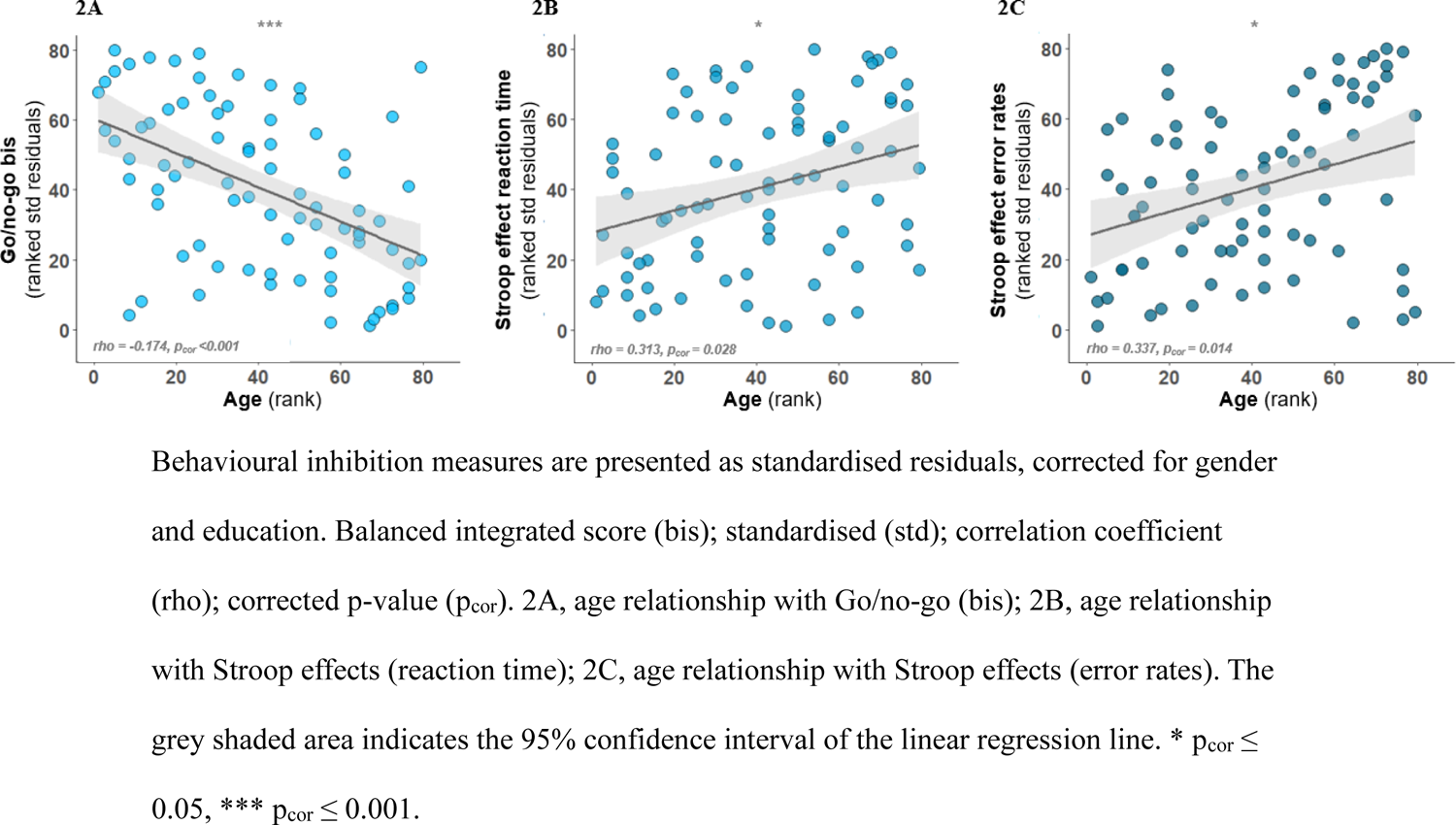
Significant Semi-Partial Correlations (Spearman’s Rank) of Age and Inhibitory Sub-Components

## Mediation Models

We conducted mediation analyses to test whether task specific inhibitory measures mediate the relationship between age and sustained attention, as measured using the *d’* and bis (see Table 4 for effect models and Figure 3 for path diagrams). Inhibitory measures were independently entered as mediators in separate models.

**Figure 3.**
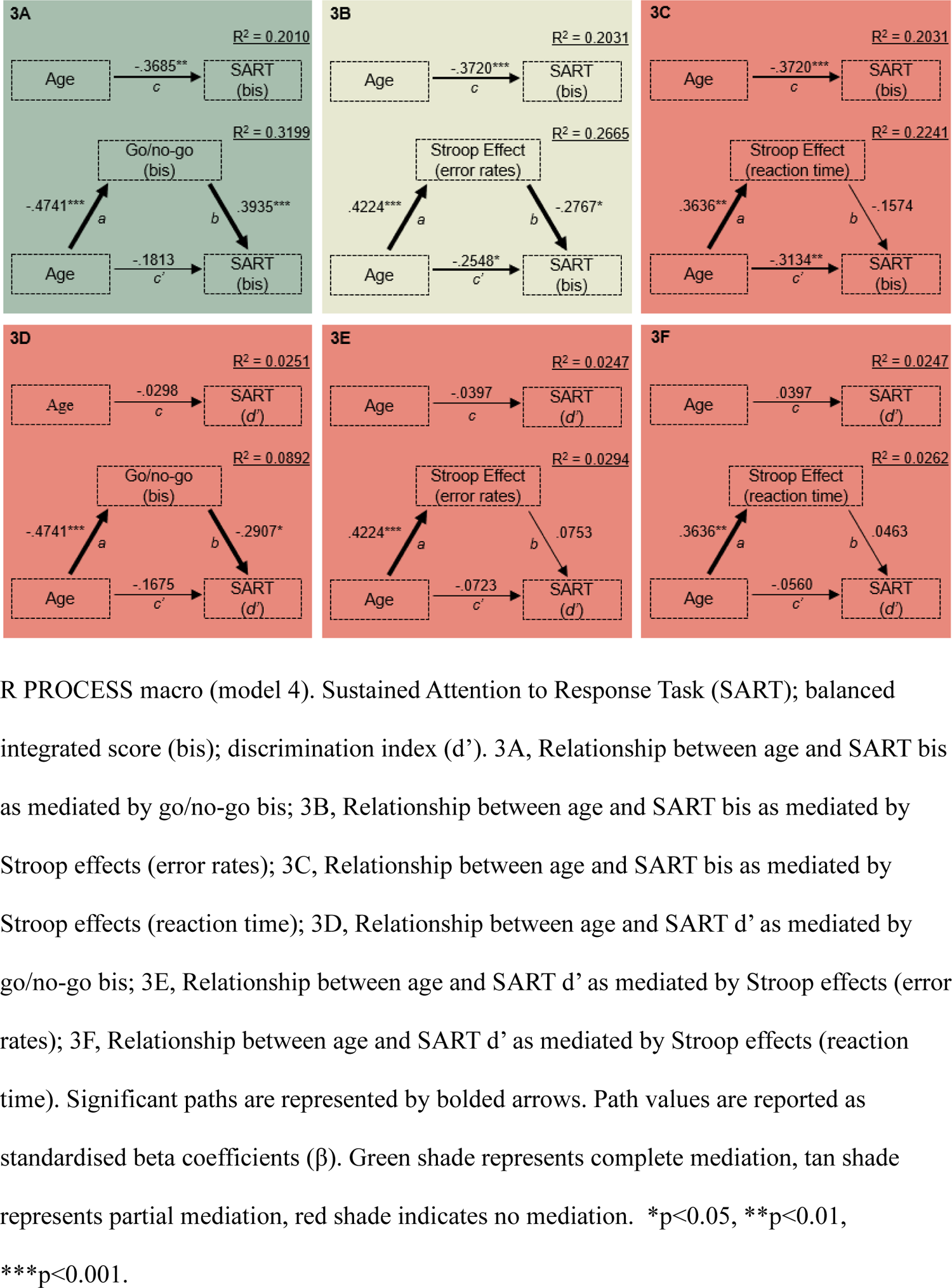
Path Diagrams for Mediation Analyses

**Table 4.**
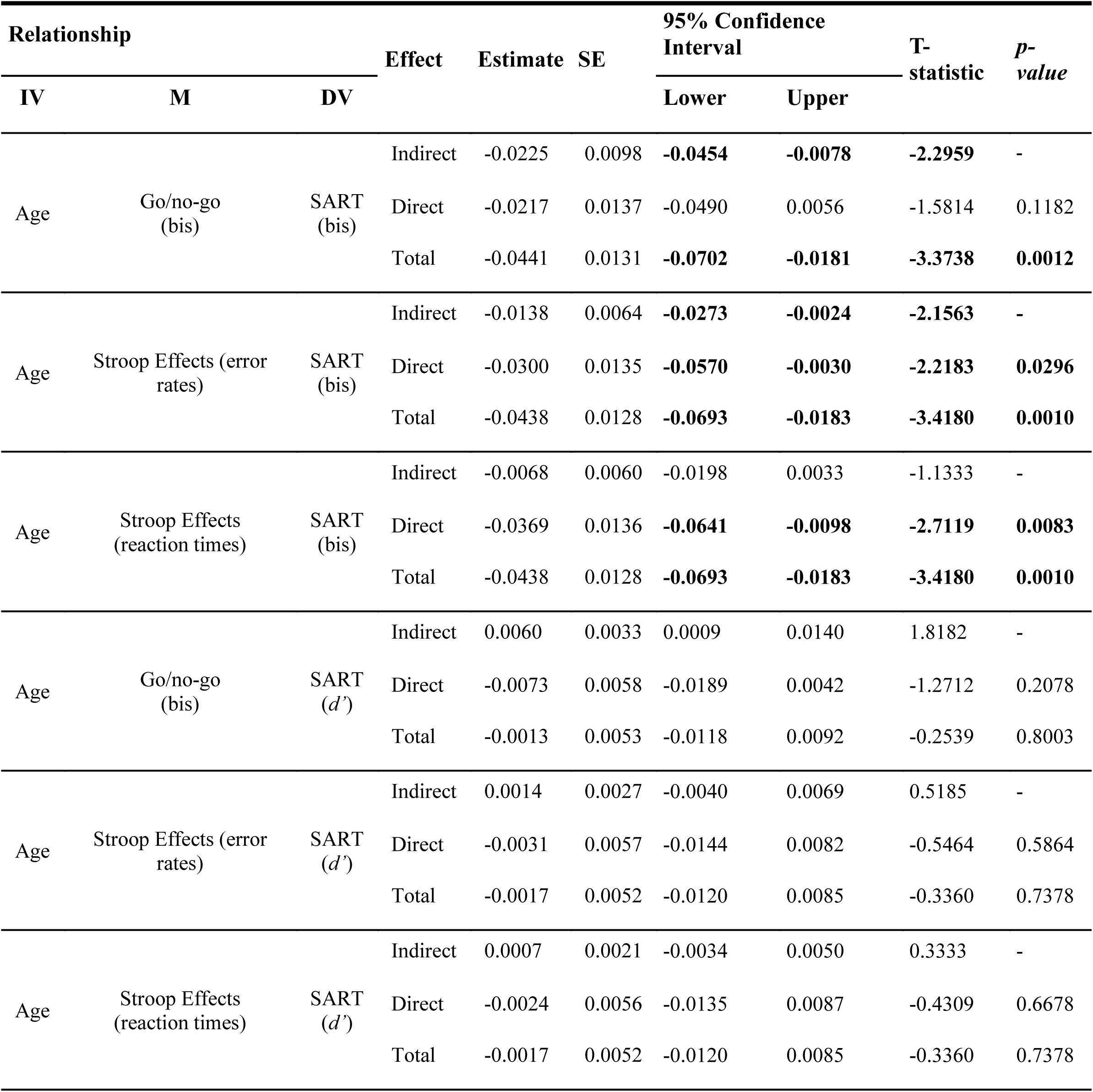
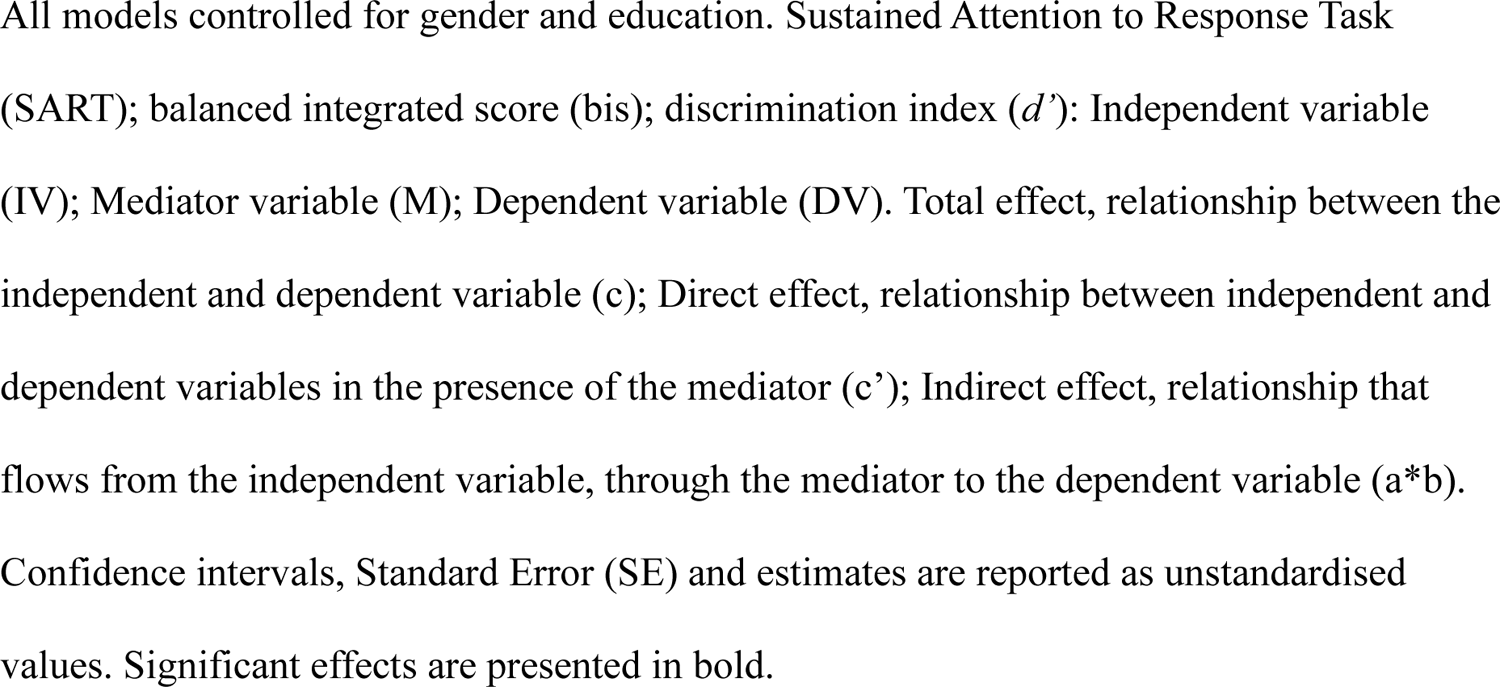
Mediation Effect Models Examining the Intervening Role of Inhibitory Functioning on the Relationship between Ageing and Sustained Attention Performance.

| Relationship |  |  | Effect | Estimate | SE | 95% Confidence Interval |  | T-statistic | p-value |
| --- | --- | --- | --- | --- | --- | --- | --- | --- | --- |
| IV | M | DV |  |  |  | Lower | Upper |  |  |
| Age | Go/no-go (bis) | SART (bis) | Indirect | -0.0225 | 0.0098 | **-0.0454** | **-0.0078** | **-2.2959** | - |
|  |  |  | Direct | -0.0217 | 0.0137 | -0.0490 | 0.0056 | -1.5814 | 0.1182 |
|  |  |  | Total | -0.0441 | 0.0131 | **-0.0702** | **-0.0181** | **-3.3738** | **0.0012** |
| Age | Stroop Effects (error rates) | SART (bis) | Indirect | -0.0138 | 0.0064 | **-0.0273** | **-0.0024** | **-2.1563** | - |
|  |  |  | Direct | -0.0300 | 0.0135 | **-0.0570** | **-0.0030** | **-2.2183** | **0.0296** |
|  |  |  | Total | -0.0438 | 0.0128 | **-0.0693** | **-0.0183** | **-3.4180** | **0.0010** |
| Age | Stroop Effects (reaction times) | SART (bis) | Indirect | -0.0068 | 0.0060 | -0.0198 | 0.0033 | -1.1333 | - |
|  |  |  | Direct | -0.0369 | 0.0136 | **-0.0641** | **-0.0098** | **-2.7119** | **0.0083** |
|  |  |  | Total | -0.0438 | 0.0128 | **-0.0693** | **-0.0183** | **-3.4180** | **0.0010** |
| Age | Go/no-go (bis) | SART (d') | Indirect | 0.0060 | 0.0033 | 0.0009 | 0.0140 | 1.8182 | - |
|  |  |  | Direct | -0.0073 | 0.0058 | -0.0189 | 0.0042 | -1.2712 | 0.2078 |
|  |  |  | Total | -0.0013 | 0.0053 | -0.0118 | 0.0092 | -0.2539 | 0.8003 |
| Age | Stroop Effects (error rates) | SART (d') | Indirect | 0.0014 | 0.0027 | -0.0040 | 0.0069 | 0.5185 | - |
|  |  |  | Direct | -0.0031 | 0.0057 | -0.0144 | 0.0082 | -0.5464 | 0.5864 |
|  |  |  | Total | -0.0017 | 0.0052 | -0.0120 | 0.0085 | -0.3360 | 0.7378 |
| Age | Stroop Effects (reaction times) | SART (d') | Indirect | 0.0007 | 0.0021 | -0.0034 | 0.0050 | 0.3333 | - |
|  |  |  | Direct | -0.0024 | 0.0056 | -0.0135 | 0.0087 | -0.4309 | 0.6678 |
|  |  |  | Total | -0.0017 | 0.0052 | -0.0120 | 0.0085 | -0.3360 | 0.7378 |

All models controlled for gender and education. Sustained Attention to Response Task (SART); balanced integrated score (bis); discrimination index ( $d'$ ): Independent variable (IV); Mediator variable (M); Dependent variable (DV). Total effect, relationship between the independent and dependent variable ( $c$ ); Direct effect, relationship between independent and dependent variables in the presence of the mediator ( $c'$ ); Indirect effect, relationship that flows from the independent variable, through the mediator to the dependent variable ( $a*b$ ). Confidence intervals, Standard Error (SE) and estimates are reported as unstandardised values. Significant effects are presented in bold.

### Mediation Analysis of Age, Response Inhibition, and Sustained Attention

When go/no-go bis was entered as a mediator of the relationship between age and SART bis (Figure 3A), significant total effects were identified (R^2^ = 0.201, *p* = 0.001), as shown in Table 4. The results revealed significant indirect effects (*t* = −2.296, CI = −0.045 to −0.008) and non-significant direct effects (p = 0.118), with the mediator accounting for an additional 11.89% of the variance (R^2^ = 0.320). In this model, age was negatively associated with go/no-go performance (such that older individuals had poorer performance), which in turn was positively associated with sustained attention performance (such that individuals with better go/no-go performance also had better sustained attention performance). In contrast, when the dependent variable was replaced with SART *d’* (instead of the SART bis measure), go/no-go performance no longer mediated the relationship between age and sustained attention (all effects were non-significant).

### Mediation Analysis of Age, Interference Inhibition, and Sustained Attention

For the model with Stroop effect (error rates) mediating the impact of age and SART bis (Figure 3B), significant total effects were identified (R^2^ = 0.203, *p* = 0.001), see Table 4. The results revealed significant indirect effects (*t* = −2.156, CI = −0.027 to −0.002) and direct effects (*p =* 0.03), with the mediator accounting for an additional 6.34% of the variance (R^2^ = 0.267). In this model, age was positively associated with Stroop performance (such that older individuals had poorer performance), which in turn was negatively associated with sustained attention performance (such that individuals with worse Stroop performance also had worse SART performance), but age also had a direct negative association with sustained attention performance (such that older individuals had poorer performance). In contrast, when the dependent variable was replaced with SART *d’* (instead of the SART bis measure), Stroop performance no longer mediated the relationship between age and sustained attention (all effects were non-significant). For the model with Stroop effect (reaction time) entered as a mediator between age and SART bis (Figure 3C), significant total effects were seen (R^2^ = 0.203, *p =* 0.001), as shown in Table 4. The results revealed a non-significant indirect effect (*t* = −1.133, CI = −0.02 to 0.003); however, a significant direct effect was observed (DR^2^ = 0.021). *p =* 0.008) Similarly, for the model with Stroop effect (reaction time) mediating the relationship between age and SART *d’*, no significant effects were identified.

## Discussion

In the present work, we tested the inhibition deficit hypothesis in a group of healthy ageing older adults (50-84 years) by investigating age-relationships with different sub-component processes of inhibition and whether inhibition can explain age-related differences in other aspects of cognition (i.e., sustained attention). Age-related declines were observed in Stroop and go/no-go performance, but not flanker performance, which suggests that age only impacts certain sub-component processes of inhibition, speaking against a *general* inhibition deficit in ageing. Taken together with the weak inter-task relationships, this task-specific age-related finding supports the independence of these inhibition sub-components, as indexed with these tasks. Importantly, we identified that go/no-go performance completely mediates the relationship between ageing and sustained attention performance, whilst Stroop effects were found to partially mediate this association. These findings are in alignment with inhibition deficit hypothesis, demonstrating that inhibitory functions may account for age-related declines in other aspects of cognitive functioning, more than ageing itself. This provides evidence that inhibition can explain differential age-effects in healthy older adults, representing an important cognitive domain to accurately maintain and monitor as we age.

### Are tasks of inhibition measuring a common construct?

In this study, we did not identify significant correlations among any of inhibition task pairs, consistent with previous work (Pettigrew & Martin, 2014; Stahl et al., 2014). The Bayesian analysis yielded mixed evidence, supporting both the null and alternate hypothesis depending on the task comparison. Notably, most inter-task correlations provided only anecdotal Bayesian evidence, lending support to the idea that these inhibition tasks are not strongly related. These findings are in general alignment with the taxonomy which separates the ability to inhibit prepotent motor responses from the ability to inhibit interfering and distracting task-irrelevant information (Stahl et al., 2014). By extension then, this indicates that these tasks, which are often described in general as, “inhibition tasks”, are likely measuring different types of inhibition. These findings demonstrate the potential pitfall of using a singular task to represent general inhibitory functioning, rather than the specific sub-component. Whilst we did not identify strong inter-task relationships in our sample of 80 individuals, we recognise that larger sample sizes might reveal stronger associations. Notably, most of the anecdotal Bayesian evidence involved correlations with the flanker task which had poor reliability (r_SB_ ≤ 0.3), while the other inhibition tasks showed moderate to good reliability estimates. This raises the possibility that the lack of strong associations amongst inhibition tasks in our study may be due to measurement reliability. Nonetheless, the weak inter-task relationships reported here and elsewhere lends support to the proposition that performance outcomes do not satisfy generalization beyond the inhibitory sub-component being measured (Rey-Mermet et al., 2018; Rush et al., 2006). Put differently, performance on a particular “inhibition” task may not inherently reflect one’s overall inhibitory ability, rather, these behavioural measures simply offer insight into one’s proficiency in a specific inhibitory process. To confound this appreciation of differences further, previous work by Raud and colleagues (2020) concluded that even tasks assumed to measure the same form of inhibition (in this case, a stop-signal and go/no-go task to measure response inhibition) seem to rely on different mechanisms which highlights their non-interchangeability. Taken together, these potentially overlooked details need to be considered when reviewing the inconsistence evidence concerning inhibition in ageing. Additionally, the utility of task paradigms with more complex manipulations has gained traction in ageing contexts (Hsieh et al., 2016; Puccioni & Vallesi, 2012; Servant & Evans, 2020), which may provide more insight into inhibitory functioning than comparably simpler effects obtained from classic tests.

### Age-related inhibitory decline is not general

Partially confirming our primary hypothesis, the results revealed a positive correlation between age and Stroop effects, aligning with previous work (Andrés et al., 2008; Fong et al., 2021). Furthermore, when a SAT score was considered, we identified a negative correlation between age and go/no-go performance. This finding reiterates the importance of accounting and/or controlling for SATs, which may well be “masking” age-inhibition relationships, leading to inadequately informed conclusions. This finding introduces yet another element of complexity to the study of inhibition, highlighting a caveat in ageing research which could be exacerbating the inconsistency, and task-dependency, of reported findings. An ageing relationship with go/no-go performance has been reported previously, however, these studies used typical dependant measures, such as reaction times and/or error rates (Nielson et al., 2002; Rey-Mermet & Gade, 2018). Here, no age relationships were identified with flanker performance or go/no-go error rates, with Bayesian evidence also supporting the null hypothesis. Of note, these semi-partial correlations were run with the contributions of gender and education regressed out of the behavioural metrics, in an effort to isolate age-effects. As such, these findings suggest that the ability to inhibit response interference and prepotent responding is vulnerable to the ageing process, whilst the ability to ignore distracting, off-task information appears to be preserved in our healthy adult cohort. These findings contrast recent work by Kouwenhoven and Machado (2024), who identified age-related deficits in both flanker and Stroop performance. As noted in the previous paragraph, the low reliability of the flanker task in our study may account this discrepancy. Moreover, our results also misalign with meta-analytic reviews by Verhaeghen (2011), who reported no age-related inhibitory decline, and Rey-Mermet and Gade (2018), who identified age-related decrements in go/no-go performance only. Although our age-inhibitory findings (i.e., age-related declines in Stroop and go/no-go, but not flanker performance) are incongruent with the task-particular outcomes of these meta-analyses, their overarching concluding criticisms, which challenge the notion of global inhibitory decline in ageing, are echoed here. The findings of the current study also speak against the hypothesis of a general inhibition deficit in ageing, but rather support the notion of inhibitory decrements with ageing among a subset of inhibitory performance measures.

### The mediating role of inhibitory sub-components

According to inhibition deficit hypothesis, inhibition can account for age-related differences in other cognitive domains (Hasher & Campbell, 2020; Hasher & Zacks, 1988). To test this, we performed a mediation analysis (controlling for gender and education) to determine whether differences in inhibition could explain age-related changes in sustained attention, over and beyond ageing itself. The results showed significant indirect effects and non-significant direct effects for go/no-go performance (SAT score) mediating the relationship between age and sustained attention performance (SAT score). This suggests that once this sub-component process of inhibition was taken into account, the relationship of age on sustained attention performance was substantially weakened, demonstrating that the ability to inhibit a prepotent response completely mediates age-related variance in attentional competencies. Thus, adequate suppression of inappropriate actions acts in service of an individuals’ capacity to be goal-directed. Further, the results demonstrated significant indirect and direct effects for Stroop effects (error rates) mediating the relationship between age and sustained attention performance (SAT score). This finding indicates that the ability to inhibit interference explains age-related differences in sustained attention alongside the direct impact of ageing itself. This implies that inhibiting interfering information may prevent attentional resources being misallocated to task-irrelevant matters, which supports the ability to sustain performance. Altogether, this asserts that specific sub-component processes of inhibition are key regulatory mechanisms subserving sustained attention in ageing, aligning with inhibition deficit hypothesis. Importantly, this also reveals a more intricate relationship between age and sustained attention which may explain disparate findings regarding direct age-attention relationships in healthy older adults (Vallesi et al., 2021), underlining the apparent role of inhibition in clarifying these associations.

The inhibition deficit hypothesis has found support previously, accounting for age-related variance in vigilance (using an auditory task) and verbal-list learning (Persad et al., 2002), as well as semantic fluency (Fong et al., 2021). To our knowledge, this is the first study to provide specific evidence that this theory extends to the domain of sustained attention (as measured using SART). That said, it is important to mention these mediation findings were again contingent on the performance measure used. When using a typical performance measure to assess sustained attention (i.e., discrimination index, *d’*), no mediation was observed, thus, these mediation effects were only identified when SATs were accounted for. The task and measure dependency of these findings, and the broader literature, emphasises the potential risk of overlooking important components of age-related change, which, depending on the task/measure used, may consequently minimise the role of inhibition in the maintenance of general and cognitive health. This highlights a nuance in ageing research, revealing the importance of targeting sub-component processes of inhibition and accounting for SATs when investigating healthy ageing, which necessitates consideration in future research.

### Strengths and Limitations

By investigating inhibition deficit hypothesis in a well-powered study of healthy older adults (n=80, 44f), utilising multiple measures of inhibition and several performance metrics, we were able to demonstrate specific age-related declines in inhibitory sub-components and the role of inhibition in mediating the effects of age on sustained attention. However, several potential limitations require consideration. This study was cross-sectional in design which impedes enquiry into causal relationships and/or temporal changes. Further, our sample of predominantly Western healthy older adults were generally very highly functioning and well-educated, which might not be truly representative of the general older adult population.

## Conclusion

The present work investigated inhibition deficit hypothesis in a group of healthy ageing older adults. We identified that only specific inhibition tasks were observed to have age-related relationships, calling into question the hypothesis of a general inhibition deficit in ageing. This result, in combination with the weak inter-task correlations, points to the non-unitary nature of commonly implemented inhibition tasks to measure “inhibition”, and the limitation of using one behavioural measure. Finally, mediation analysis revealed that specific inhibitory sub-components account for differential age-effects in other cognitive domains (here, sustained attention), over and beyond ageing itself, via an indirect path.

Though we provide partial support for inhibition deficit hypothesis with sub-component processes of inhibition, it is important to emphasise the task and measure dependency of these highly nuanced findings, underscoring the pitfall of conflating these sub-components into a generalised concept of inhibition. This is coupled with the need to attenuate speed-accuracy trade-offs in ageing research, which likely mask age-effects. Bearing these cautions in mind, it is reasonable to conclude that inhibitory sub-components and SAT scores must be considered in ageing contexts to reveal associations which may be otherwise obscured or overlooked.

