## Supplementary material for "Age-related Inhibitory Decline: Examining Inhibition Sub-Components and their Impact on Sustained Attention in Healthy Ageing": supp material

**Supplemental Material**

**Table S1**

*Median reaction time and accuracy data for the flanker and Stroop tasks.*

|  |  | **N** | **Min-max** | **Mean(±SD)** |
| --- | --- | --- | --- | --- |
| **Flanker** | Reaction time (CT) | 80 | 538 - 1246 | 766.1(±146.35) |
|  | Reaction time (ICT) | 80 | 577.5 – 1160 | 788.04(±137.10) |
|  | Accuracy (CT) | 80 | 84.53 - 100 | 97.25(±3.56) |
|  | Accuracy (ICT) | 80 | 83.42 - 100 | 96.56(±3.3) |
|  | Reaction time (CT) | 80 | 661 - 1169 | 869.50(±110.18) |
| **Stroop** | Reaction time (ICT) | 80 | 682 - 1428 | 960.84(±144.94) |
|  | Accuracy (CT) | 80 | 94.67 - 100 | 99.44(±1.14) |
|  | Accuracy (ICT) | 80 | 80.67 - 100 | 97.51(±4.37) |

Congruent trial (CT); Incongruent trial (ICT)

**Table S2**

*Mean reaction time and accuracy data for the go/no-go task.*

|  |  | **N** | **Min-max** | **Mean(±SD)** |
| --- | --- | --- | --- | --- |
| **Go/no-go** | Reaction time | 79 | 292.38 – 554.63 | 426.35(±61.03) |
|  | Accuracy | 79 | 60 - 100 | 90.44(±8.58) |

**Table S3**

*Mean reaction time and probability of hits and false alarms for the SART.*

|  |  | **N** | **Min-max** | **Mean(±SD)** |
| --- | --- | --- | --- | --- |
| **SART** | Reaction time | 78 | 126.03 – 522.38 | 224.53(±68.10) |
|  | False alarms; commission errors | 78 | 0.10 – 0.98 | 0.56(±0.21) |
|  | Hit rate; omission errors | 78 | 0.73 – 0.99 | 0.93(±0.05) |

Sustained Attention to Response Task (SART). False alarms and hit rate are log linear corrected values, with mean performances ranges on a scale from 0 to 1.

**Table S4**

*Estimated internal consistency of behavioural measures using a permutation-based splithalf approach with 5000 random splits*

| **Behavioural Measure** | **n** | **Spearman-Brown corrected estimates** | **95% CI** |
| --- | --- | --- | --- |
| Stroop Effect (accuracy) | 80 | 0.80 | 0.74 – 0.91 |
| Stroop Effect (reaction time) | 80 | 0.78 | 0.69 – 0.85 |
| Flanker Effect (accuracy) | 80 | 0.30 | -0.07 – 0.55 |
| Flanker Effect (reaction time) | 80 | -0.14 | -0.39 – 0.19 |
| Go/no-go (accuracy) | 79 | 0.76 | 0.66 – 0.84 |
| Go/no-go (reaction time) | 79 | 0.63 | 0.50 – 0.73 |
| SART (accuracy) | 78 | 0.82 | 0.76 – 0.87 |
| SART (reaction time) | 78 | 0.96 | 0.94 – 0.97 |

Sustained Attention to Response Task (SART). The splithalf package in R (Parsons, 2021) does not have the bis or d’ as a scoring option, thus, these SART reliability estimates are for the constituent accuracy and reaction time data.

Parsons, S. (2021). Splithalf: Robust estimates of split half reliability. *Journal of Open Source Software*, *6*(60), 3041.
